# New SARS-CoV-2 lineages could evade CD8+ T-cells response

**DOI:** 10.1101/2021.03.09.434584

**Authors:** Marco Antonio M. Pretti, Rômulo G. Galvani, Alessandro S Farias, Mariana Boroni

## Abstract

**Background:** Many SARS-CoV-2 variants of concern have emerged since the Covid-19 outburst, notably the lineages detected in the UK, South Africa, and Brazil. Their increased transmissibility and higher viral load put them in the spotlight. Much has been investigated on the ability of those new variants to evade antibody recognition. However, not enough attention has been given to pre-existing and induced SARS-CoV-2-specific CD8+ T cell responses during the natural course of infection by new lineages.

**Methods:** In this work, we investigated the SARS-CoV-2-specific CD8+ T cell epitopes from the main variants of concern and the potential of associated mutations to trigger or hinder CD8+ T-cells response. We also estimated the population’s coverage of these different lineages, considering peptide binding predictions to class I HLA alleles from 29 countries to investigate differences in the fraction of individuals expected to respond to a given epitope set from new and previous lineages.

**Results:** We observed a lower populational coverage for 20B/S.484K (P.2 lineage) in contrast to an increased coverage found for 20H/501Y.V2 (B.1.351 Lineage) and 20J/501Y.V3 (P.1 lineage) compared to a reference lineage. Moreover, mutations such as Spike N501Y and Nucleocapsid T205I were predicted to have an overall higher affinity through HLA-I than the reference sequence.

**Conclusions:** In summary, the data in this work provided evidence for the existence of potentially immunogenic and conserved epitopes across new SARS-CoV-2 variants, but also highlights the reduced populational’s coverage for the Brazilian lineage P.2, suggesting its potential to evade from CD8+ T-cell responses. Our results also may guide efforts to characterize and validate relevant peptides to trigger CD8+ T-cell responses, and design new universal T-cell-inducing vaccine candidates that minimize detrimental effects of viral diversification and at the same time induce responses to a broad human population.

## Background

Since the Covid-19 outburst, researchers are struggling in the entire world to better understand why the disease manifests in many different ways, generating a vast range of symptoms. During the last months, new SARS-CoV-2 lineages were detected in the United Kingdom, South Africa, and Brazil (1). Some of these new mutations are of global interest since they seem to increase SARS-CoV-2 transmission (2,3). Most variants of concern have multiple mutations in the S gene (4) which codes for immunodominant peptides (5). The emergence of these new lineages raised concerns about the reinfection of convalescent individuals as well as the effectiveness of available vaccines. Indeed, mRNA vaccines-elicited antibodies showed a reduced plasma neutralizing activity against new lineages (6). Consistently, Brazilian P.1 lineage seems to escape from neutralizing antibodies generated against previously circulating variants of SARS-CoV-2 as well as inactivated virus Covid-19 vaccine-elicited antibodies (7).

Interestingly, although neutralizing antibody titers and the memory B cell response are short-lived in almost all human coronavirus infections, there is neither a high number of reinfections nor a large number of recurrences of serious illness (SARS, MERS, or Covid19) (8,9). In this context, adaptive cellular response plays an important role in protective immunity against SARS-CoV-2 (8,10). Many studies have evidenced the role of T-cell response to SARS-CoV-2 (5,11,12). SARS-CoV-2-specific CD8+ T cells from individuals with milder Covid-19 have higher perforin and granzyme activity than convalescent ones, showing an increased CD8+ T effector phenotype during the acute mild disease (11). Of note, the cytotoxic activity is also increased in CD4+ T cells responding to SARS-CoV-2 (13).

Moreover, convalescent individuals showed naturally occurring CD4+ and CD8+ memory T cells specific for Spike (S), Nucleocapsid (N), Membrane, and some non-structural ORF proteins leading to the production of single or multiple proinflammatory cytokines (IL-2, TNF-ɑ, and IFN-γ) (5,11,14–16). These results highlight the importance of pre-existing and induced SARS-CoV-2-specific CD8+ and CD4+ T cell responses for immune protection in mild SARS-CoV-2 infection. Since then, studies suggested the inclusion of specific peptides to elicit T cell response in vaccine design and *in silico* analysis of SARS-CoV-2 proteins has detected candidate epitopes specific for protective T CD4+ or CD8+ and B responses with low risks of allergy or autoimmunity (17–20). In fact, approved vaccines induce T cell response in addition to the production of neutralizing antibodies for immunization against SARS-CoV-2 (21–23).

The quality and amplitude of adaptive immunity are highly dependent on the Human Leukocyte Antigen (HLA) complex-mediated presentation of epitopes to T-cells. Previous works have already evidenced sets of SARS-CoV-2 peptides more likely to be presented through HLA molecules to CD8+ T-cells (24–26). In a recent study, our group performed a computational approach to map populational coverage for SARS-CoV-2-derived peptides based on class I HLAs, which suggested a protective role of a higher S/N coverage ratio (26). Since most of the new SARS-CoV-2 variants are located in the proteins S and N, we decided to investigate whether mutations in these regions could modify peptide presentation, antigenic coverage, and impact in T cell response.

## Methods

### SARS-CoV-2 lineages

BetaCoV/Wuhan/IPBCAMS-WH-01/2019 genome assembly (GenBank: MT019529.1) was downloaded and used as a wild-type reference (REF) sequence. Non-synonymous Single-Nucleotide Variants (nsSNVs) and deletions in S and N genes of four SARS-CoV-2 upcoming variants of interest (VOI) (20I/501Y.V1, 20H/501Y.V2, 20J/501Y.V3, and 20B/S.484K) were retrieved from the online database covariants.org (4). A list of S and N mutations analyzed here can be found in Table S1.

### Binding and antigenicity predictions

A matrix containing the allele frequencies of 297 class I HLA-A and -B alleles from 37 countries was obtained (26) considering the cumulative allele frequency of those alleles as close to 0.9 as possible for each country. Binding predictions were performed as previously described (26) and peptides classified as Strong (SB) and Weak Binders (WB) were kept for the analysis. Importantly, residues from positions −10 to +10 in relation to the mutated amino acid (aa) were used to predict 8 to 11-mer binders. Only peptides that were distinct between each VOI and the REF were considered in the analysis. Peptide antigenicity was predicted to all supported HLA alleles among the 297 using PRIME (27) with default parameters by providing a list of all binders predicted by netCTLpan (28) and filtering for TAP and Cle scores as previously described (26). Scores with zero values were excluded since they mean no HLA presentation.

### Population coverage

Predicted SB and WB class I HLA-I:peptides pairs derived from SARS-CoV-2 VOI and REF were used to calculate population coverage, using the IEDB Population coverage software (29). The software was downloaded on 02/02/2021 and ran locally with default parameters for binders exclusive to REF or VOI lineages. In this analysis, 29 countries were included (Austria, Brazil, Bulgaria, China, Croatia, Czech Republic, England, France, Germany, Indonesia, Israel, Italy, Japan, Malaysia, Mexico, Morocco, Oman, Poland, Portugal, Romania, Russia, Senegal, Spain, Thailand, Tunisia, Ireland Northern, South Africa, United States, and India). Among the initial list of 37 countries used in (26), 9 were removed due to insufficient data (cumulative coverage of less than 50% for one epitope hit). In addition, India was included in the list.

### Statistical and data analysis

The analysis was conducted on the R environment v4.0, and we used the R packages Biostrings v2.56 for peptide sequence manipulation and alignment, ggmsa v0.0.5 for visualization, and ggpubr v0.3.0 for statistical analysis. Wilcoxon test was used for mean comparisons unless stated otherwise. A p-value < 0.05 was used as a threshold for mean comparisons except for the antigenicity score where a p<0.01 was used instead.

## Results

Focused on evaluating the binding profile to class I HLA of some of the SARS-CoV-2 variants, 26 nsSNVs and two deletions located within the genes coding for proteins S and N were obtained (4). The mutations represent the genetic variability within these two proteins of four VOI circulating worldwide. We have originated a set of 8 to 11-mer peptides from 21-length aa sequences comprising each variant, within a window of 10 residues before and after the variant whenever possible (Figure S1).

Following binding predictions for the 8 to 11-mer peptides to the 297 class I HLA-A and -B molecules, 27 peptides shared between REF and respectives VOI were filtered out and 230 unique peptides remained in the analysis. Of note, none of the remained viral peptides matched the human proteome. Those 230 peptides generated 6,320 unique HLA-I:peptide combinations. Comparing the number of HLA-I alleles predicted to bind REF and VOI-derived peptides, we observe that peptides derived from three variants in S (69-70del, A570D, Y144del) lack the ability to bind to the HLA alleles set (Table S2, Figure 1). Importantly, five out of 21 remaining nsSNVs in protein S generated fewer HLA-I:peptide pairs when compared to the REF sequence, and two of which (D614G and T1027I) generated less than half the number of binders compared to the respective REF sequences. Notably, the nsSNV N501Y generated 10 times more binders than the REF sequence. All nsSNVs from protein N generated more binders compared to the REF sequences and, surprisingly, neither the nsSNV S235F nor REF sequence generated any binders. The absence of peptide ligands derived from nsSNVs/deletions could contribute to the diminishing cellular immunity whilst a higher number of epitopes may favor a T cytotoxic response. Studying the relationship between antigen presentation and different mutations across VOI may shed light on population susceptibility to specific viral lineages.

**Figure 1.**
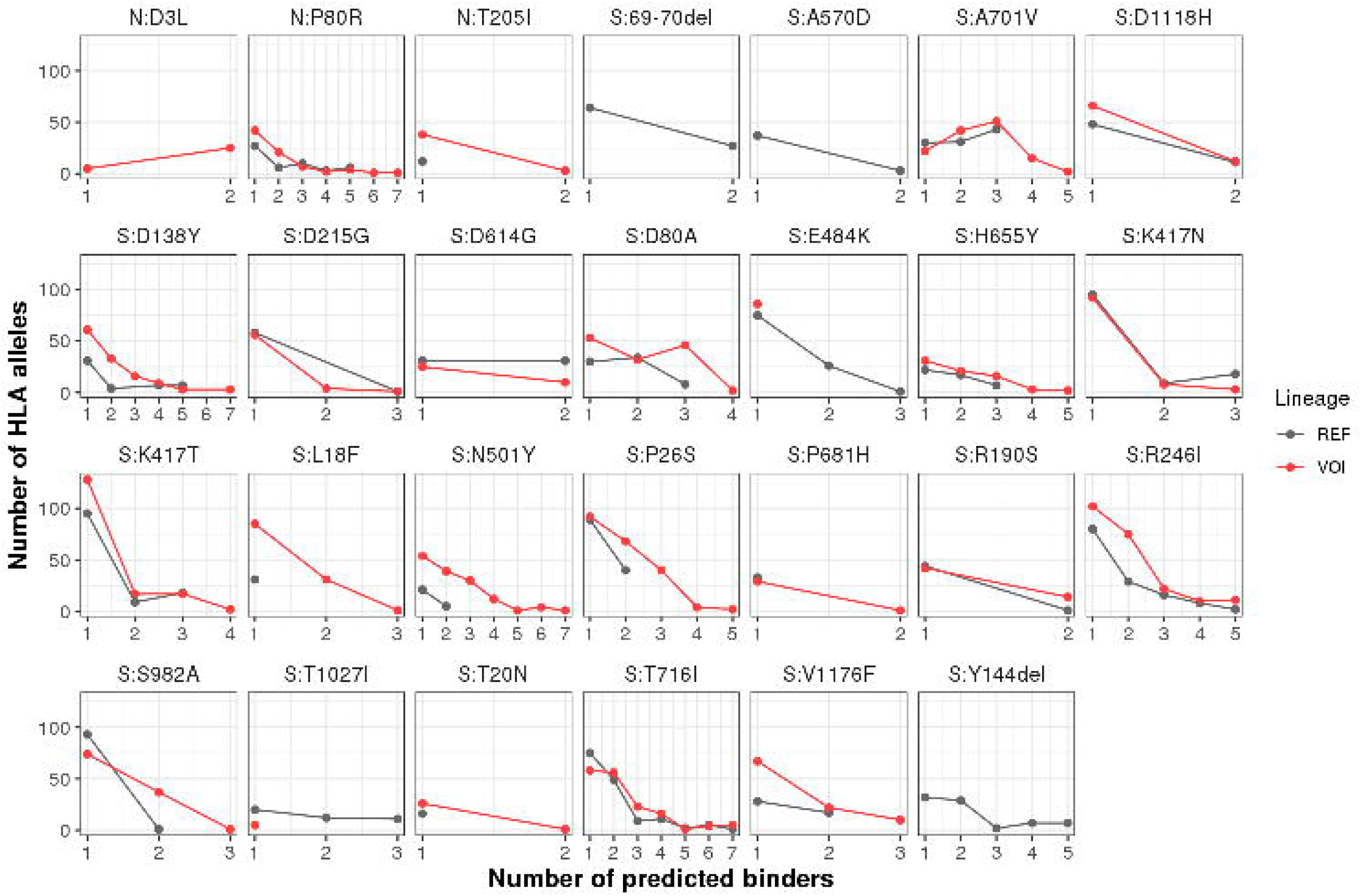
Number of unique HLA-I:peptide pairs for REF and VOI-derived peptides. Binding predictions considering 297 class I HLA-A and -B alleles were performed on VOI and REF aa sequences. For each nsSNV/deletion, the distribution of the number of HLA alleles per number of predicted binders is depicted for VOI (Red) and REF (Black) peptides. The nsSNV N:S235F did not generate any binder (Table S2).

Although we verified an increased number of potentially presented peptides derived from all mutations in protein N and the majority in protein S (15/24), lower affinities of variant peptides to the HLA-I molecule could counterbalance their elevated number. Therefore, we compared the HLA-affinities between VOI and REF-derived peptides (Figure S2). A lower %Rank indicates peptides more likely to bind a given allele when compared with a set of natural binder peptides. A significantly increased affinity was observed for a pool of six peptides derived from the nsSNVs T205I, D80A, H655Y, N501Y, P26S, and T716I while the ones derived from E484K, K417N, and T1027I exhibited lower affinity compared to the respective REF-derived peptides. Of note, for the VOI 20I/501Y.V1, new mutations resulted in epitopes with significantly increased affinities in protein S, while for epitopes derived from 20B/S.484K, a significant decrease in affinity was observed when compared to the REF lineage (Figure S2).

Recently, a few studies released information about the *in vitro* reactivity of SARS-CoV-2 epitope-specific T cell responses (5,30,31). We then accessed data from Tarke and collaborators (5) to check whether our binding predictions matched the *in vitro* data. We identified an overlap of 15 REF-derived peptides with their set of 191 peptides derived from proteins S or N, corresponding to 18% of REF-derived peptides identified by our predictions. When considering only the REF derived-epitopes predicted to bind to the 10 HLAs with more *in vitro* testing, this number increases to 23%. Of note, the REF-derived peptides investigated by Tarke and collaborators that induce T cell response fall into the following mutations found in the VOIs: T716I P80R, 69-70del, A701V, D215G, D614G, D80A, H655Y, K417T, K417N, P26S, P681H, S982A, T1027I, V1176F, and Y144del. Importantly, the VOI-derived peptides were not considered in this analysis as they were not individually tested *in vitro (30)*.

Interested in investigating the antigenicity of SARS-CoV-2 variants, we compared REF and VOI-derived peptides using a predictor that considers TCR recognition of HLA-restricted epitopes (27). We obtained 16,019 non-zero peptide:HLA combinations and observed that nsSNVs P80R, A701V, E484K, H655Y, K417N, K417T, L18F, R246I, S982A, and T716I had greater antigenicity scores in regard to the REF sequence (Figure S3). In contrast, nsSNVs T205I, D1118H, D614G, P26S, and Y144del showed lower antigenicity scores. Considering all nsSNVs associated with each VOI, an increased antigenicity was observed for 20H/501Y.V2 (p-value < 0.0001) and 20J/501Y.V3 (p-value <0.001) lineages (Figure 2). We then associated those results with our previous biding predictions (Figure1, Figure S2) considering only peptides with significant differences between REF and VOI lineages. Interestingly, mutation-derived peptides that were predicted as more antigenic could be classified into two groups, one predicted to be less presented and the second having higher presentation scores. Regarding the less antigenic ones, all of them were predicted to be better presented (Table 1). Of note, 20H/501Y.V2 had three nsSNVs with lower antigenicity scores despite better epitope presentation. Overall, our data indicate that some REF-derived peptides are in the immunodominant region. Thus, changes in this specific region may culminate in the escape of the virus from the protective response of T cells, either in convalescent or vaccinated individuals.

**Figure 2.**
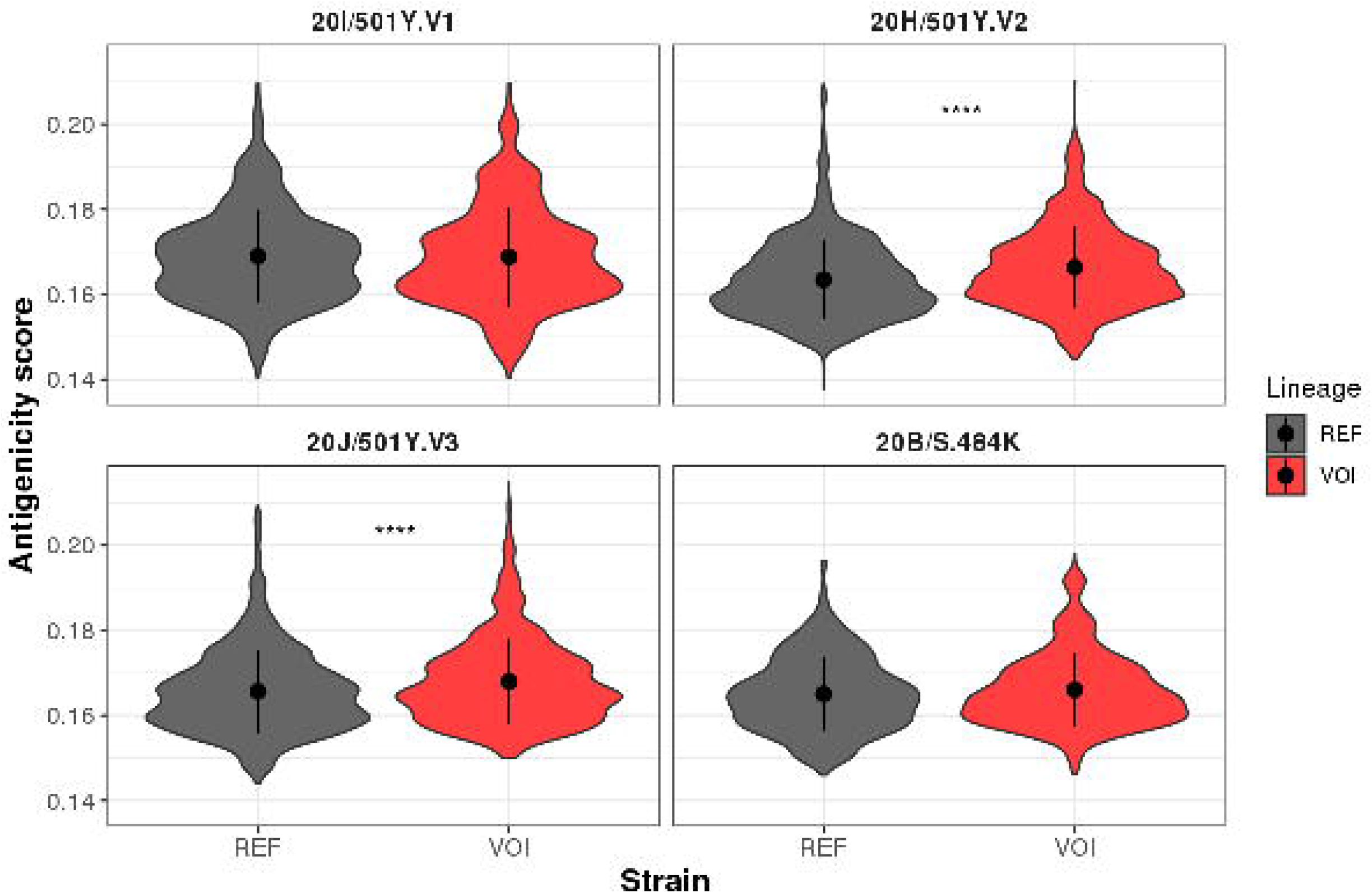
Overall antigenicity of SARS-CoV-2 VOI. Predictions were obtained using the list of 297 HLA alleles to the PRIME algorithm that calculated the likelihood of a peptide being presented through the HLA-I and triggering a T-cell response. Non-zero scores for all peptide:HLA combinations are shown. All nsSNVs were aggregated according to the respective lineage they belong to. *** p<0.001; ****p<0.0001.

**Table 1.** Comparison between presentation and antigenicity scores between REF and VOI lineages.

| Group | Mutations | Predicted HLA presentation | Antigenicity score | Associated VOI |
| --- | --- | --- | --- | --- |
| Reduced antigenicity despite better presentation | T205I, P26S | VOI > REF | REF > VOI | 20H/501Y.V2<br>20J/501Y.V3 |
| Increased antigenicity and better presentation | H655Y, T716I | VOI > REF | VOI > REF | 20J/501Y.V3<br>20I/501Y.V1 |
| Increased antigenicity and worse presentation | E484K, K417N | REF > VOI | VOI > REF | 20H/501Y.V2,<br>20J/501Y.V3,<br>20B/S.484K |

Despite the differences in binding profile and antigenicity, the HLA alleles-presenting SARS-CoV-2 binder peptides are not necessarily equally distributed across different countries. Therefore, we investigated how different populations are predicted to be covered for the four VOI compared to the REF lineage using HLA frequency data from 29 countries. In this specific analysis, we considered the allele frequency of the class I HLA alleles in each analyzed population and the number of S and N-derived peptides (epitope hits) they are likely to present (Figure 3). We observed a significant difference (p<0.0001, Wilcoxon) when comparing the area under the curve of VOI and REF coverages for 20H/501Y.V2 (coverage ratio=1.32), for 20J/501Y.V3 (coverage ratio=1.27), and for 20B/S.484K (coverage ratio=0.74) lineages but not for 20I/501Y.V1 (ratio=1.02, p=0.58). Together, these findings indicate remarkable differences in antigen coverage among the lineages and the absence of HLA-I presentation for particular nsSNVs that could lead to evasion from T-cell responses.

**Figure 3.**
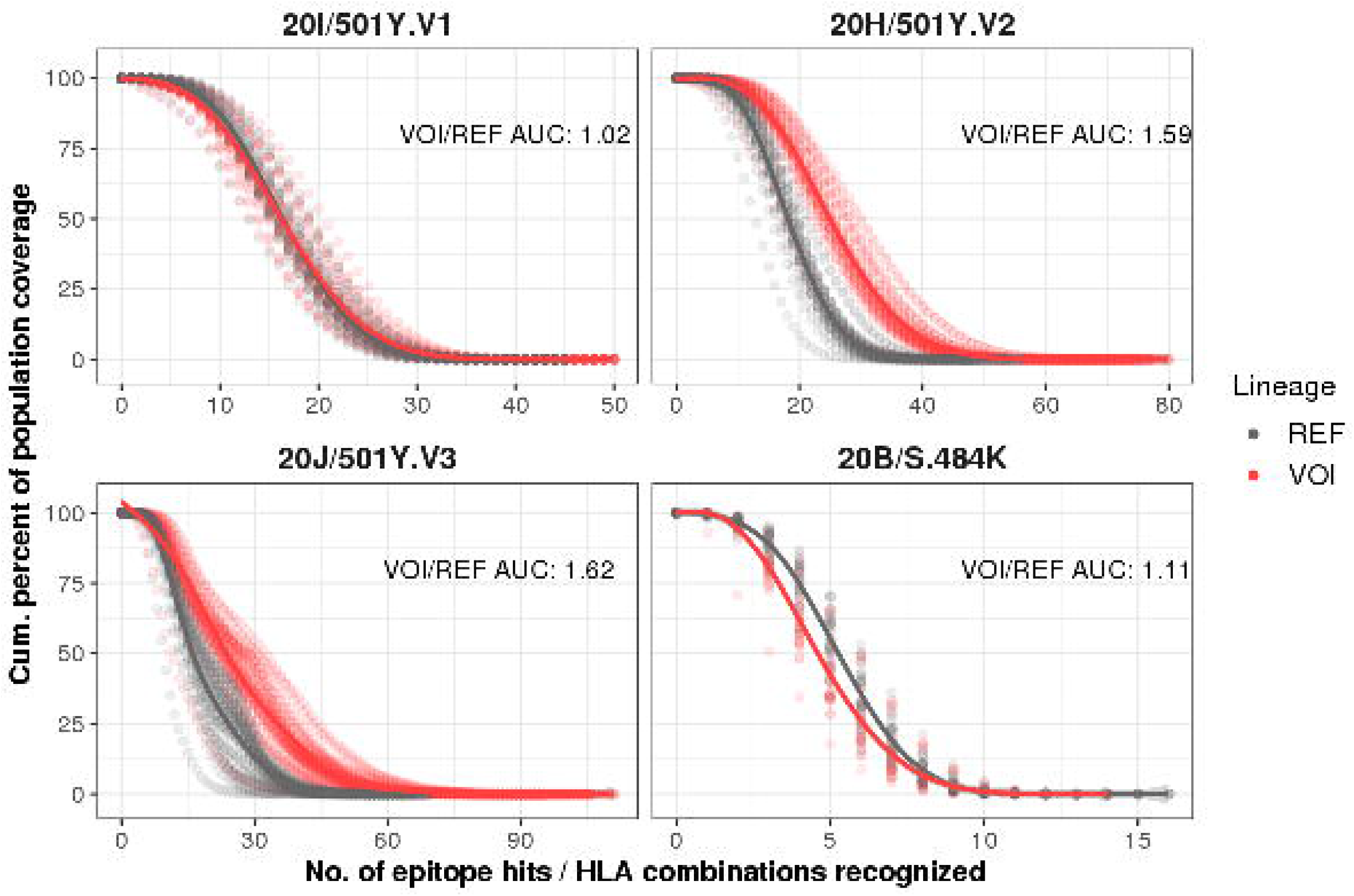
Populational coverage for the four SARS-CoV-2 VOI compared to the REF sequence. Population coverage analyses were based on 297 HLA-A and -B alleles using allele frequency data from 29 countries. Each panel represents a different VOI (red) compared with the REF (gray). Tendency lines were drawn connecting the VOI and REF dots. The VOI/REF area under the curve (AUC) ratio is depicted.

## Discussion

The outburst of new SARS-CoV-2 variants capable of faster spreading than previous lineages is a major concern for health systems worldwide. Four of them bear nsSNV that confer more transmissibility than other circulating coronaviruses (2,3), named 20I/501Y.V1 (B.1.1.7 Lineage), emerged in Britain and detected in over 70 countries; 20H/501Y.V2 (B.1.351 Lineage), emerged in South Africa, and two that emerged in Brazil, 20J/501Y.V3 (P.1 Lineage) and 20B/S.484K (P.2 Lineage) (32). Cytotoxic CD8+ T cells are key for immune protection against viral infections, and recent works have shown the importance of cross-reactive and induced SARS-CoV-2-specific CD8+ T cell responses to immune protection (5,31,33). Given the role of SARS-CoV-2-specific CD8+ T cells for Covid-19 resolution, we investigated the likelihood of four SARS-CoV-2 lineages to generate new epitopes capable of binding a set of class I HLA alleles representing countries from Europe, Americas, Africa, and Asia to ultimately infer T-cell mediated immunity.

Limited data exist on the magnitude and extension of CD8+ T cell responses associated with the new circulating SARS-CoV-2 variants. Mutations leading to new putative epitopes, or either the lack of conserved ones should be investigated for their potential to generate exacerbated responses or evade the immune response. We observed that mutations such as 69-70del in the gene coding for protein S did not generate HLA-I binders and such deletions were associated with antibody escape in a recent work (34). The number of putative HLA ligands derived from REF or VOI sequences gives us an idea of whether there is material for T-cell recognition since a higher number of peptides increases the probability of a robust and effective T cell response.

Another important aspect is whether the mutations-derived peptides are more or less immunogenic and likely to trigger the T-cell response. For instance, E484K and K417N substitutions have overall less affinity to class I HLA molecules than the respective REF peptides. Although not mandatory to trigger a T-cell response, tighter binding to HLA-I groove facilitates T-cell response (35,36). On the other hand, both were predicted to have higher antigenicity than the REF peptides which may indicate their potential of promoting immune system evasion. Importantly, these nsSNVs are located on the Receptor Binding Domain (RBD), the region responsible for the entry of the virus into the cell (37). This finding is in line with other reports that identified evasion from antibodies associated with E484K and K417N nsSNVs (37,38). On the other hand, 501Y and 655Y substitutions have a higher affinity to the HLA alleles than the REF peptides but, similarly to E484K, are able to escape from antibody recognition (39,40). Considering the increased affinity to the HLA, N501Y and H655Y substitutions may be good targets to trigger CD8+ T-cell responses. Despite no difference in affinity, the nsSNV D614G that has been related to increased transmissibility (2,3) displayed half the number of potential binders than 614D. Interestingly, the substitutions T205I in the protein N and P26S in protein S had an increased affinity to the HLA despite lower antigenicities, which may point to a positive selection of less immunogenic peptides. In addition, our previous work suggested presentation of S-derived peptides rather than N-derived peptides could be beneficial for the individual. Based on that, the fact that 20I/501Y.V1 lineage potentially presents more N-derived epitopes could be a major concern to the protective response in convalescent and vaccinated individuals. In fact, all nsSNVs on the protein N, e.g. D3L, P80R, and T205I, generated more potential epitopes in comparison to the REF peptides. To date, there are no studies associating the nsSNVs in protein S such as D80A, P618H, T1027, and T716I with increased transmissibility or immune evasion, since most studies focused on nsSNVs located on the RBD. Comparing the predictions and the peptides tested *in vitro* (5), we noticed that the overlap was higher when considering the most frequent alleles. This is expected since *in vitro* testing does not usually consider lower frequency HLAs as it is the case of *in silico* predictions. Moreover, the overlap with peptides known to trigger T-cell responses indicates that losing them may have negative effects in recognizing SARS-CoV-2 epitopes. In fact, a recent work described that SARS-CoV-2 mutations may help evading from CD8+ T-cell responses (41).

The overall susceptibility to new SARS-CoV-2 lineages is a major concern. We have used a populational coverage strategy to investigate whether the predicted ability of a given population to recognize and present the viral epitopes is impacted by the new mutations in the VOIs. While the investigated countries exhibited higher coverage for the lineages 20H/501Y.V2 and 20J/501Y.V3, a lower coverage was observed for the 20B/S.484K lineage, firstly identified in patients in Rio de Janeiro, Brazil (42). A case of reinfection was reported for this lineage (43) and to this date, it was already detected in Europe, North and South America (32). Of note, Spain was the country less covered for all lineages except for 20B/S.484K when considering both REF and VOI coverages. Although more studies are still necessary, the data indicate less presentation of peptides from 20B/S.484K lineage, and may suggest its ability to evade T cell response, justifying its high spreadability, being detected in multiple other countries including England, Singapore, the USA, Norway, Argentina, Denmark, Ireland, and Canada (42). Importantly, in January 2021, Brazil was considered the worst country in the management of the Covid-19 pandemic, according to a study by the Lowy Institute (44). The combination of low testing, low number of Covid-19 vaccination doses administered and ineffective protection measures that culminate in a high number of infected individuals will make Brazil the main source of new SARS-CoV-2 lineages in the coming months. Finally, besides investigating the populational T-cells response to SARS-CoV-2 new lineages, the ability of the available vaccines to protect against those new lineages should be further explored (33,45). The reduction or even lack of epitopes derived from protein S in the VOI could impact the effectiveness of some available vaccines. On the other hand, being aware of the binding profile of new peptide variants may guide vaccine development efforts by ranking conserved epitopes of interest, thus reducing or eliminating the need to select new epitopes from emerging lineages.

## Conclusions

Overall, our *in silico* analysis of antigenic coverage revealed a lessen HLA coverage associated with the 20B/S.484K lineage among human populations, suggesting that the CD8+ T response may be affected when compared to the wild SARS-CoV-2 lineage, probably enhancing the risk of pathogen escape in naturally exposed individuals.

## Abbreviations

aa: amino acid
HLA: Human Leukocyte Antigen
IFN-γ: Interferon gamma
IL-2: Interleukin-2
MERS: Middle east respiratory syndrome
N: Nucleocapsid
nsSNVs: Non-synonymous single-nucleotide variants
RBD: Receptor Binding Domain
REF: reference lineage
S: Spike
SARS: Severe acute respiratory syndrome
SARS-CoV-2: Severe acute respiratory syndrome coronavirus 2
SB: Strong Binder
TCR: T-cell receptor
TNF-ɑ: Tumor necrosis factor
VOI: variant of interest
WB: Weak Binder

## Ethics approval and consent to participate

Not applicable.

## Availability of data and materials

The sequence of SARS-CoV-2 can be found at the GenBank with the ID: MT019529.1. The list of HLA-I alleles is available in a previous publication (26). All software used in this work is free for academic use.

## Authors’ contributions

MB, MP, and RG contributed to the design and conception of the study. MB and ASF contributed to important scientific discussions. MP and RG performed all the analyses and wrote the first draft of the manuscript. MB was responsible for the final approval of the submitted version. All the authors contributed to the article and approved the submitted version.

## Acknowledgments

The authors would like to thank the Bioinformatics Core Facility (INCA-RJ) for their support.

## Supplementary Figure Legends

**Figure S1**. Global alignment of the 21-length aa sequence for each variant of four SARS-CoV-2 VOI to the REF sequence. Variants are surrounded by ten aa residues on each side, except by the nsSNV N:D3L, in which the variant falls at the beginning of the protein. Dashes (-) represent deletions (del) within the sequence.

**Figure S2.** Binding affinities of VOI and respective REF-derived peptides. Binding affinities predictions (%Rank score) considered 297 class I HLA-A and -B alleles on 8 to 11-mer peptides derived from VOI (Red) and respective REF sequences (Gray). The lower the %Rank score the higher the likelihood of a peptide to bind a given HLA molecule. The nsSNV 235F did not generate any binder (Table S2). *p< 0.05, **p<0.01, ***p<0.001, ****p<0.0001

**Figure S3.** Predicted antigenicity of SARS-CoV-2 nsSNVs. For each nsSNV, unique HLA-I:peptide pairs were used to predict their overall TCR antigenicity. Only p values below 0.01 were considered significant. **p<0.01; ***p<0.001; ****p<0.0001.

