## Supplemental Table 1 for "New SARS-CoV-2 lineages could evade CD8+ T-cells response"

Table S1. List of SARS-CoV-2 nsSNV used in the analysis.

| **VOI** | **nsSNV in protein S** | **nsSNV in protein N** |
| --- | --- | --- |
| 20I/501Y.V1 | N501Y, D614G, 69-70del, Y144del, A570D, P681H, T716I, S982A, D1118H | D3L, S235F |
| 20H/501Y.V2 | L18F, K417N, E484K, N501Y, D614G, D80A, D215G, R246I, A701V | T205I |
| 20J/501Y.V3 | L18F, K417T, E484K, N501Y, D614G, T20N, P26S, D138Y, R190S, H655Y,T1027I | P80R |
| 20B/S.484K | E484K, D614G, V1176F | - |
