## Supplemental Table 2 for "New SARS-CoV-2 lineages could evade CD8+ T-cells response"

Table S2. Number of unique HLA-I:peptide pairs derived from deletions and nsSNVs in proteins N and S considering the four VOI and REF sequences.

| **nsSNV / deletion** | **HLA-I:peptide pairs** | | | |
| --- | --- | --- | --- | --- |
|  | **WB** | | **SB** | |
|  | **REF-derived peptides** | **VOI-derived peptides** | **REF-derived peptides** | **VOI-derived peptides** |
| N:D3L | 0 | 50 | 0 | 5 |
| N:P80R | 77 | 113 | 34 | 33 |
| N:S235F | 0 | 0 | 0 | 0 |
| N:T205I | 12 | 30 | 0 | 14 |
| S:69-70del | 90 | 0 | 28 | 0 |
| S:A570D | 20 | 0 | 23 | 0 |
| S:A701V | 155 | 258 | 66 | 71 |
| S:D1118H | 59 | 70 | 11 | 20 |
| S:D138Y | 77 | 207 | 25 | 40 |
| S:D215G | 38 | 43 | 23 | 24 |
| S:D614G | 74 | 31 | 19 | 14 |
| S:D80A | 84 | 152 | 38 | 111 |
| S:E484K | 103 | 75 | 27 | 11 |
| S:H655Y | 75 | 123 | 2 | 20 |
| S:K417N | 114 | 99 | 53 | 18 |
| S:K417T | 114 | 167 | 53 | 54 |
| S:L18F | 24 | 132 | 7 | 18 |
| S:N501Y | 31 | 199 | 0 | 107 |
| S:P26S | 123 | 240 | 46 | 134 |
| S:P681H | 25 | 26 | 8 | 5 |
| S:R190S | 32 | 47 | 14 | 23 |
| S:R246I | 181 | 311 | 47 | 102 |
| S:S982A | 55 | 101 | 40 | 50 |
| S:T1027I | 42 | 5 | 35 | 0 |
| S:T20N | 9 | 19 | 7 | 9 |
| S:T716I | 208 | 230 | 83 | 137 |
| S:V1176F | 50 | 100 | 12 | 41 |
| S:Y144del | 108 | 0 | 51 | 0 |
