## Supplementary figures and images for "New SARS-CoV-2 lineages could evade CD8+ T-cells response"

### Supplemental Figure 1

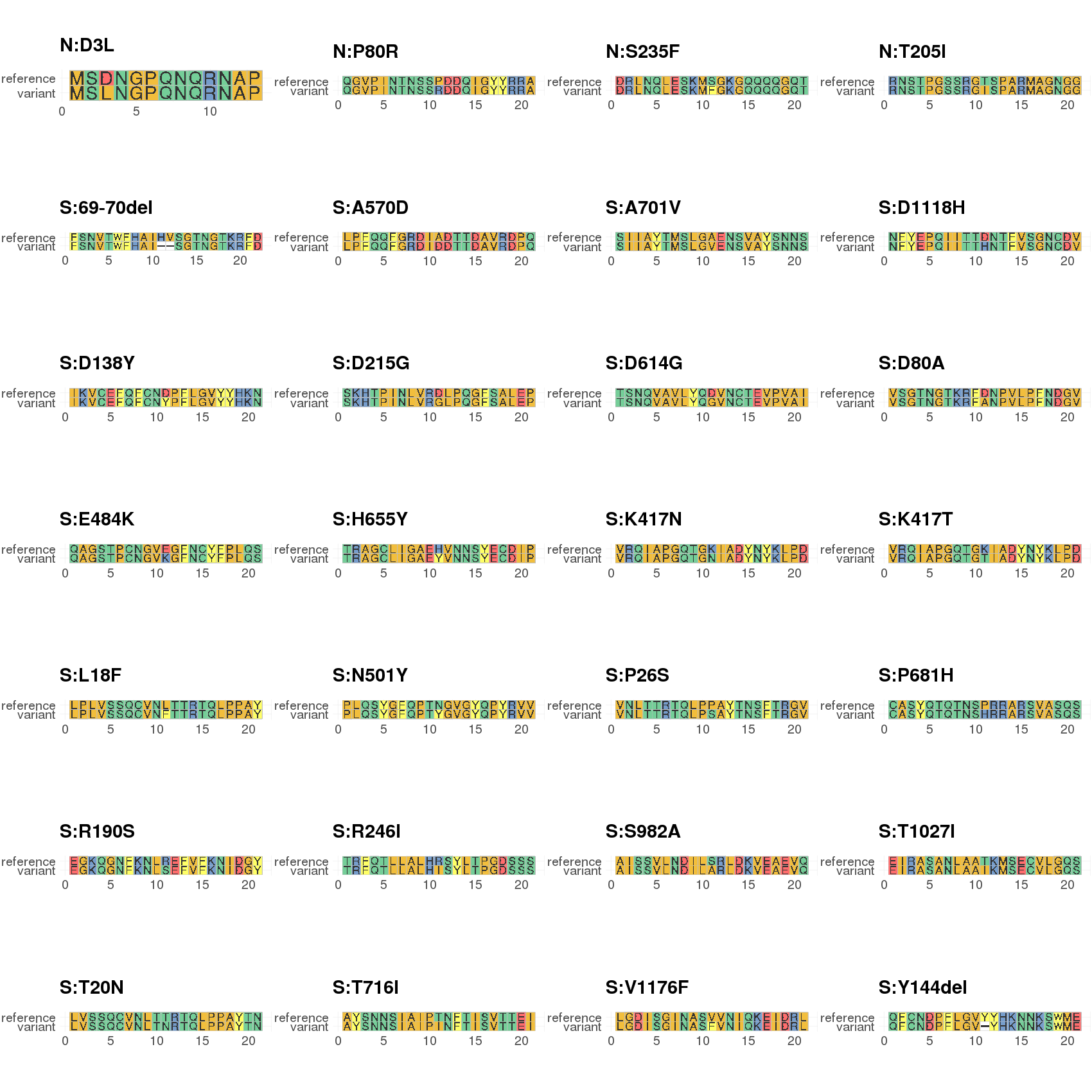

### Supplemental Figure 2

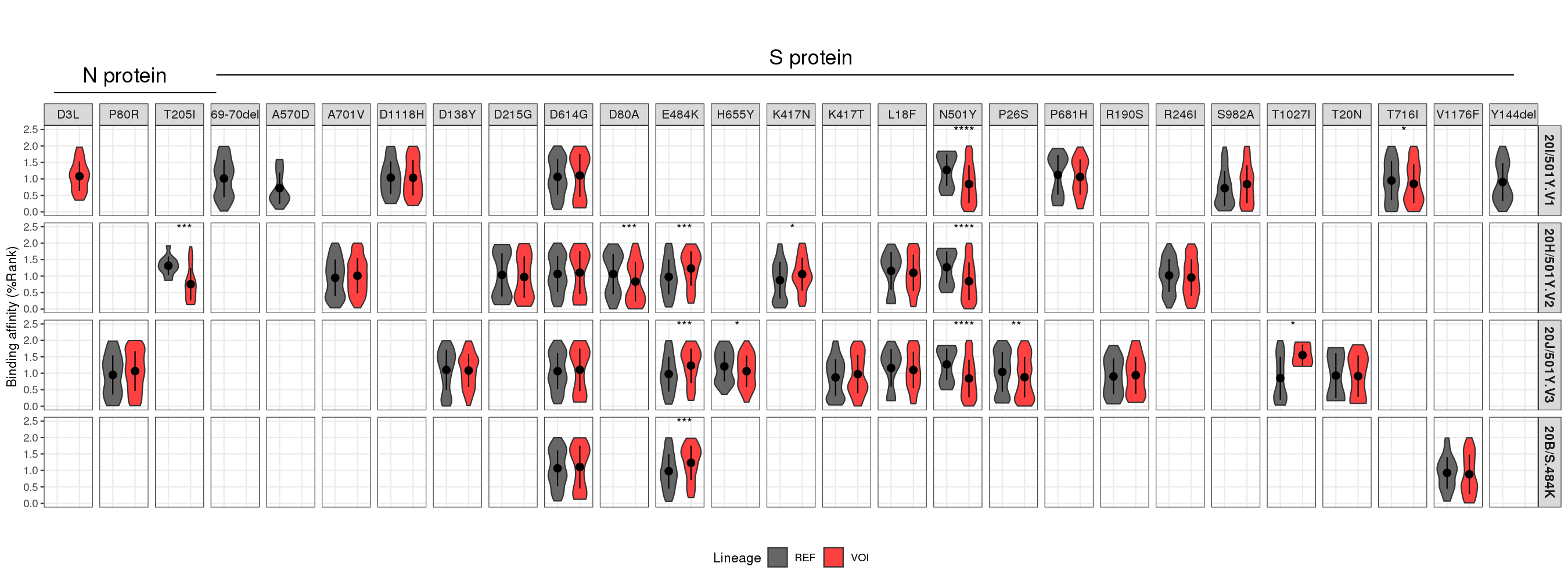

### Supplemental Figure 3

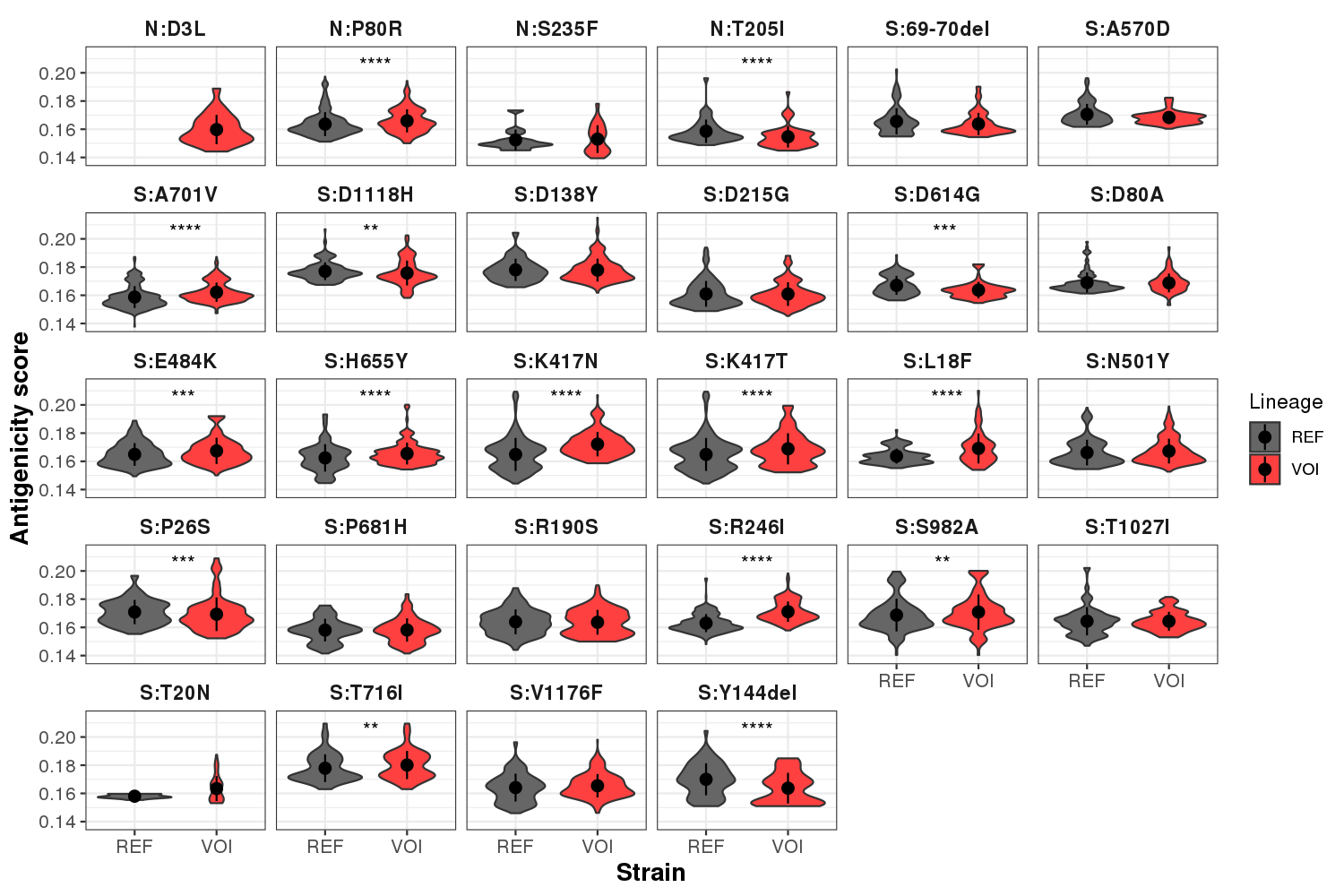
